# Distraction and cognitive control independently impact parietal and prefrontal response to pain

**DOI:** 10.1101/2022.06.20.496774

**Authors:** Nicolas Silvestrini, Corrado Corradi-Dell’Acqua

## Abstract

Previous studies found that distracting someone through a challenging activity leads to hypoalgesia, an effect mediated by parietal and prefrontal processes. Other studies suggest that challenging activities affect the ability to regulate one’s aching experiences, due to partially-common neural substrate between cognitive control and pain at the level the medial prefrontal cortex. We investigated effects of distraction and cognitive control on pain by delivering noxious stimulations during or after a Stroop paradigm (requiring high cognitive load) or a neutral condition. We found less intense and unpleasant subjective pain ratings during (compared to after) task execution. This hypoalgesia was associated with enhanced activity at the level of the dorsolateral prefrontal cortex and the posterior parietal cortex, which also showed negative connectivity with the insula. Furthermore, multivariate pattern analysis revealed that distraction altered the neural response to pain, by making it more similar to that associated with previous Stroop tasks. All these effects were independent of the nature of the task which, instead, led to a localized neural modulation around the anterior cingulate cortex. Overall, our study underscores the role played by two facets of human executive functions, which exert independent influence in the neural response of pain.

## Introduction

Previous studies established that pain engages and interacts with high-level executive functions (Tracey and Mantyh, 2007; Bushnell *et al*., 2013; Wiech, 2016). For instance, distracting someone through a concurrent activity might reduce one’s sensitivity to pain (Petrovic *et al*., 2000; Tracey *et al*., 2002; Valet *et al*., 2004; Buhle and Wager, 2010). At the neural level, distraction hypoalgesia has been associated with decreased activity in regions relevant for pain processing, like somatosensory and insular cortices (Petrovic *et al*., 2000; Valet *et al*., 2004; Seminowicz and Davis, 2007b), but also with enhanced signal within the prefrontal cortex and periaqueductal gray (Petrovic *et al*., 2000; Tracey *et al*., 2002; Valet *et al*., 2004). These effects have been interpreted consistently with the neurocognitive model of attention to pain according to which bottom-up nociceptive inputs can engage one’s attention in competition with top-down processes, such as prefrontal and parietal mechanisms for maintaining attentional load and the processing of goal-relevant stimuli (Legrain *et al*., 2009).

Despite the overall consensus, several considerations put into question the role played by distraction on pain sensitivity. For instance, some studies failed to replicate the effect (McCaul *et al*., 1992; Duker *et al*., 1999) or, at least, limit its efficacy to a subgroup of individuals (Keogh *et al*., 2000; Seminowicz *et al*., 2004; Nouwen *et al*., 2006; Seminowicz and Davis, 2007a). Furthermore, distraction is often manipulated by delivering pain *During* the execution of highly demanding tasks (e.g., Stroop, N-Back) which, however, exert extensively individual control/regulatory processes by asking to select an appropriate response among multiple competing information. Cognitive control might influence pain experience independently of distraction, as suggested by studies describing changes in the sensitivity to pain delivered *After* a Stroop task (Silvestrini and Rainville, 2013; Hoegh *et al*., 2019; Silvestrini *et al*., 2020) compared to an easier activity such as counting neutral words. Furthermore, pain and cognitive control disclose distinct but co-localized neural networks (Kragel *et al*., 2018; Silvestrini *et al*., 2020), which are integrated at the level of the medial prefrontal cortex (Shackman *et al*., 2011; Silvestrini *et al*., 2020), partly reminiscently of the neural structures held to mediate distraction hypoalgesia. In this view, it is unclear whether (and to which extent) effects of distraction on pain are partly confounded by those of cognitive control. Partial evidence in favour of the independence of the two processes arises by studies modulating parametrically the cognitive load of the distracting task, who failed to find a consequent linear influence on pain sensitivity (Hodes *et al*., 1990; McCaul *et al*., 1992; Seminowicz and Davis, 2007b). However, to our knowledge, the effects of distraction and cognitive control on the behavioural and neural response to pain have never been investigated independently.

In this study, we administered noxious thermal stimulations *During* or *After* (factor TIMING) an interfering Stroop (with high control load) or an easier paradigm requiring counting neutral words (factor TASK), whilst recording pain-related ratings and neural response through fMRI. We predicted typical effects of distraction when pain was administered *During* a task, with associated modulations at the level of prefrontal-parietal structures. Moreover, we tested effects of cognitive control when comparing the Stroop Interference *vs*. the easier condition. The critical question, however, is whether TIMING and TASK influence pain experience independently, or whether instead they interact with one another.

## Methods

### Participants and Design

Twenty-nine participants (11 males, mean age = 23.86 ± 4.75 SD) were recruited by announcement at the University of Geneva. Participants were free of self-reported acute or chronic pain, and cardiovascular, neurological, or psychological disease. They signed an informed consent prior to the experiment. This research was conducted in accordance with the Declaration of Helsinki and was approved by the local ethical committee.

### Painful Thermal Stimulation

A computer controlled thermal stimulator with a 25 x 50 mm fluid-cooled Peltier probe (MSA Thermotest, Somedic, Hörby, Sweden) delivered painful thermal stimulations. The baseline temperature was set to 36°C and each stimulation lasted 16 sec (3 sec of temperature increase, 10 sec of plateau, and 3 sec of temperature decrease). The temperature of the painful stimulations was adjusted for each participant using a calibration procedure.

### fMRI Data Acquisition

A Siemens Trio 3-T whole-body scanner was used to acquire both T1-weighted anatomical images and gradient-echo planar T2*-weighted MRI images with blood oxygenation level dependent (BOLD) contrast. Structural images were acquired with a T1-weighted 3D sequence (repetition time = 1900 msec, inversion time = 900 msec, echo time = 2.27 msec, 1 × 1 × 1 mm voxel size). The functional sequence was a trajectory-based reconstruction sequence with a repetition time (TR) of 2100 msec, an echo time (TE) of 30 msec, a flip angle of 90 degrees, in-plane resolution 64x64 voxels (voxel size 3 x 3 mm), 32 slices, a slice thickness of 3 mm, and no gap between slices.

### Experimental Paradigm

Before the scanning session, participants were seated in front of a computer. The probe of the thermal stimulator was installed on their right leg in the lower part of the shinbone and the individual painful temperature was identified through staircase procedure (Silvestrini *et al*., 2020; Silvestrini and Corradi-Dell’Acqua, 2022). We applied three series of ascending and descending thermal stimulations. Participants rated the unpleasantness of each stimulation with a visual analogue scale (VAS) ranging from 0 (*Not unpleasant at all*) to 100 (*The most unpleasant pain imaginable*). After an initial increase by steps of 2°C, the temperature started decreasing when participants rated pain unpleasantness as more than 70 and increased again when pain unpleasantness was lower than 50. The last series of ascending and descending stimulations included steps of 1°C. Based on participants’ ratings, we selected a temperature that elicited a pain unpleasantness response of about 70 (temperature: *M* = 47.07, *SD* = 1.83, range = 42-51). The selected temperature remained constant throughout the main experiment.

Then, participants were engaged in a practice session for a Stroop paradigm from previous research (Silvestrini and Rainville, 2013; Silvestrini *et al*., 2020; Riontino *et al*., 2022). They were exposed to sets of one-to-four identical words presented vertically on the screen in capital letters (font Verdana size 24). They were asked to report the number of words displayed (regardless of their meaning) as quickly/accurately as possible. Moreover, participants were asked not to blur their vision preventing them to count effortlessly the words. In the interference condition, the words used was one of “one”, “two”, “three”, and “four” (in French, “un”, “deux”, “trois”, “quatre”), so that it was always inconsistent with the correct response. In the neutral condition, we used instead words of matched length which were not held to elicit any interference: “year”, “sheet”, “table”, and “skittle” (in French, “an”, “drap”, “table”, and “quille”). Trials started with a fixation cross (750ms) followed by the words which stayed on the screen until a response was delivered (but never more than 1250ms). Participants then received feedback about their proficiency.

Subsequently, participants were installed on the scanner bed, where they underwent two functional runs of 21 min each, separated by the anatomical scan (see Figure 1, for the design structure). Both scanning runs began with a painful thermal stimulation used as a baseline pain induction. After the stimulation, participants rated subjective pain intensity and unpleasantness by moving with their fingers a cursor displayed on the screen over two VAS, ranging from *No pain/Not unpleasant at all* (0) to *The most intense/unpleasant pain imaginable* (100). Then participants performed four blocks of the Stroop task. However, differently from the practice session, the feedback provided was uninformative of one’s performance. The feedback appeared for 2 sec minus participants’ reaction time, assuring that all trials had the same length (3.4 sec). Inter-trial intervals varied randomly (750-1500 ms). In the *During* condition, the blocks included 36 trials (2 min 30 sec) and a thermal stimulation was administered at the 32^nd^ trial while the participants were performing the remaining four trials. In the *After* condition, the blocks included 32 trials (2 min) and a stimulation was administered right after the end of the task. Each run included the four possible blocks (*During/Interference*, *After/Interference*, *During/Neutral*, and *After/Neutral*), which were separated by a short break of 12 sec. The order of the blocks was counterbalanced within the runs. At the end of the task and stimulation, participants rated pain and unpleasantness associated with the thermal event (total rating time: 24sec) and one item of perceived task difficulty (“How difficult was it for you to achieve the task successfully?”; *Not at all* [0] to *Very difficult* [10]).

**Figure 1.**
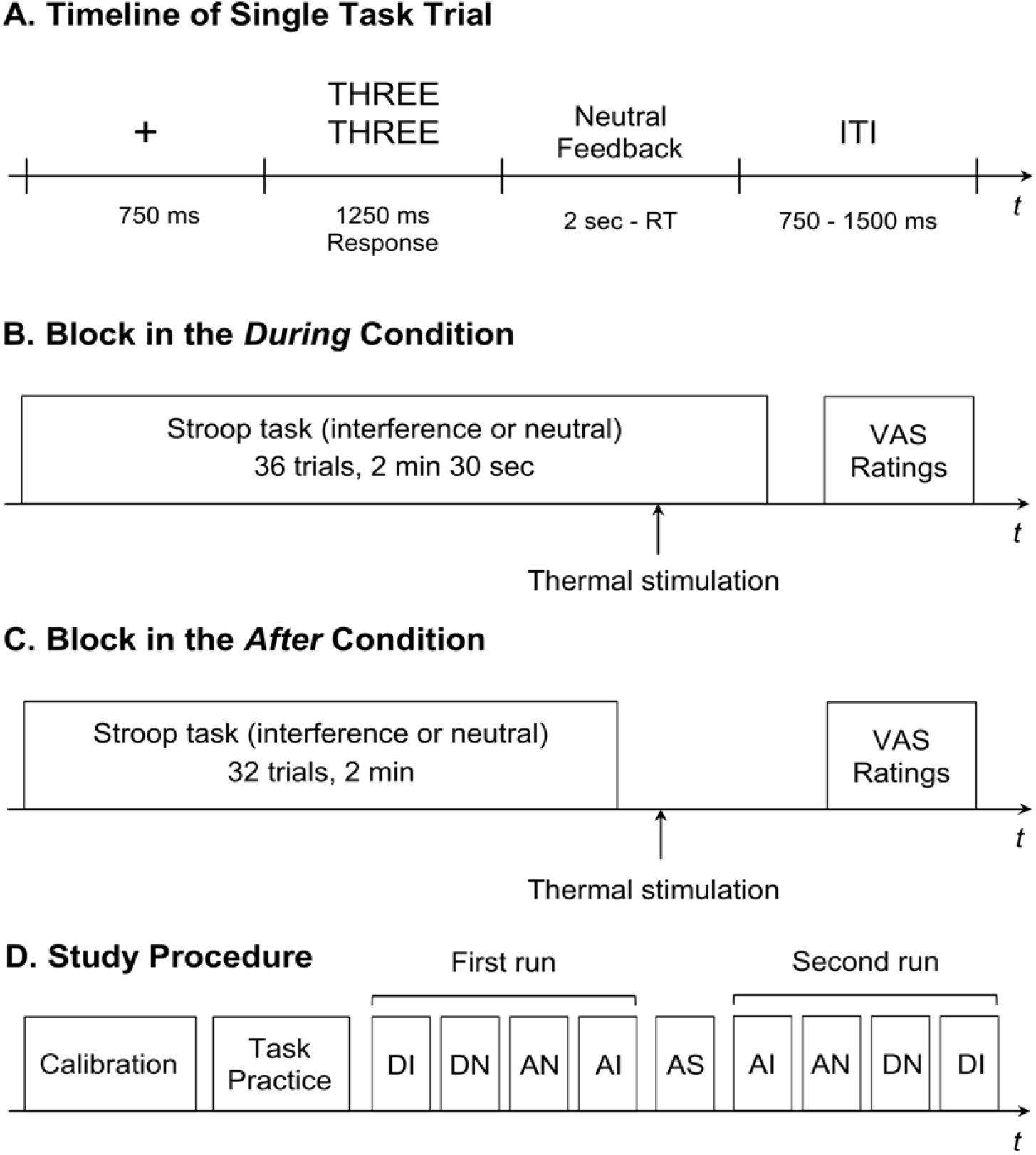
Timeline of a single task trial (Panel A), overview of experimental blocks in the During (Panel B) and the After (Panel C) conditions, and overview of the study protocol (Panel D). Neutral feedback: “Response recorded” or “Please answer more quickly” in case of no response, ITI: Intertrial interval, VAS: Visual analogue scales, AS: Anatomical scan, DI: During-Interference, DN: During-Neutral, AN: After-Neutral, AI: After-Interference. The order of the experimental conditions was counterbalanced across subjects.

The experiment was administered using Eprime 2.0 Software (Psychology Software Tools Inc., Pittsburgh, PA, USA). Images were displayed on a computer monitor back projected onto a screen and viewed on a mirror placed on the head coil.

### Data Processing

#### Behavioural Data

For each subject and condition, the average Accuracy and median correct Reaction Times from Stroop trials were analysed with a paired-sample t-test probing differences between Interference *vs*. Neutral TASK conditions. As for participants’ ratings, average Intensity and Unpleasantness values were analysed with Repeated Measures ANOVA, with TIMING (*During vs. After*) and TASK (*Interference vs. Neutral*) as within-subject factors. The analysis was run using R 4.0.4 software (http://cran.r-project.org).

#### Imaging processing

##### Preprocessing

Statistical analysis was performed using the SPM12 software (http://www.fil.ion.ucl.ac.uk/spm/), exploiting the preprocessing pipeline from the CONN 21 toolbox (https://web.conn-toolbox.org/, Whitfield-Gabrieli and Nieto-Castanon, 2012). For each subject, functional images were realigned, unwrapped, and slice-time corrected. The Artifact Detection Tools, embedded in the CONN toolbox, were then used for identification of outlier scans in terms of excessive subject motion and signal intensity spikes. Finally, the images were then normalized to a template based on 152 brains from the Montreal Neurological Institute (MNI) with a voxel-size resolution of 2 x 2 x 2 mm, and smoothed by convolution with an 8 mm full-width at half-maximum Gaussian kernel.

##### First-Level analysis

Preprocessed images from each task were analysed using the General Linear Model (GLM) framework implemented in SPM. Our key event of interest were pain trials. Hence, for each functional run, we modelled the onset of each thermal stimulation with a boxcar function of 10 sec duration starting the moment in which thermode reached the plateau temperature. Thermal events were specified separately according to the associated experimental condition (*During/Interference*, *After/Interference*, *During/Neutral*, and *After/Neutral*). Additionally, the baseline stimulation, conducted at the beginning of each functional run, was modelled as a separate vector. In addition to the main analysis, we also looked at the neural responses evoked by the Stroop task, where each of the two conditions (*Interference* and *Neutral*) was delivered through ∼2 minutes blocks. Although optimal for testing cognitive control after-effects on pain (Silvestrini and Rainville, 2013; Silvestrini *et al*., 2020; Riontino *et al*., 2022), such long block structure would make the analysis of task-related activity vulnerable to low-frequency confounds and poorly accords with default filtering in SPM (128 sec). To circumvent this, we analysed specific Stroop trials in an event-related fashion as a delta function (without distinguishing between *Interference* and *Neutral* condition) and specifying an additional predictor in which trials Reaction Times were modulated parametrically. This parametric predictor was our measure of interest for Stroop-related activity, as it should capture both differences between *Interference* and *Neutral* conditions (*Interference* trials take on average longer, see results) but also inter-trial fluctuations in performance and, as such, should be less vulnerable to signal low-frequency noise/drifts. Supplementary Information provides additional analyses on an independent dataset (Verstynen, 2014), showing that such approach represents a reliable test for neural responses of Stroop demands. Overall, we specified 7 predictors for each functional run (4 main thermal stimulations, 1 baseline thermal stimulation, 1 Stroop, 1 Reaction Times parametrical modulation), which were convolved with a canonical hemodynamic response function and associated with their first order temporal derivative. To account for movement-related variance, and other sources of noise, we included the 6 differential movement parameters from the realignment (x, y, and z translations and pitch, roll, and yaw rotations), and dummy variables signalling outlier scans (from the ART toolbox) as covariates of no interest. Low-frequency signal drifts were filtered using a cut-off period of 128 sec, and serial correlation in the neural signal were accounted through first-order autoregressive model AR(1).

##### Second-level Analyses

Functional contrasts, testing differential parameter estimates images associated with one experimental condition *vs.* the other were then analysed through a second level one-sample t-test using random-effect analysis. Effects were identified as significant only if they exceeded a FWE cluster-level correction for multiple comparisons at the whole brain (Friston *et al*., 1993), with an underlying voxel-level threshold corresponding to *p* < 0.001 (uncorrected).

##### Functional Connectivity

Preprocessed images were denoised through the default pipeline in CONN toolbox to remove components in the neural signal which were related to (1) white matter and cerebro-spinal fluid signal (first 15 principal components), (2) estimated subject movement parameters (from preprocessing), (3) the presence of outlier scans (estimated through the ART toolbox during preprocessing), and (4) task-related BOLD signals (in our case, pain-evoked activity and stroop trials). Data was also band-pass filtered (0.008-0.09 Hz) to account both for slow-frequency fluctuations and well as physiological and residual movement artefacts. Subsequently, denoised data was analyzed through ROI-to-ROI connectivity using generalized psychophysiological interaction (gPPI) (McLaren *et al*., 2012). For this analysis we considered the ROIs from Brainnetome parcellation of the human brain (Fan *et al*., 2016). This allows for a whole-brain fine-grain exploration of the regions involved without any a priori assumption on the implicated network. In particular, we focused on 224 ROIs comprehending the whole atlas, except for the regions 189-210 corresponding to the occipital cortex (Fan et al., 2016, Table 1) which were outside our range of interest. gPPI is a task-dependent connectivity analysis computing how strongly the BOLD response time course of two regions are coupled in a given condition (in our case, thermal stimulations). This is implemented though a regression model, whereby the time course of target ROIs is regressed against that of a seed ROI multiplied with the psychological variable. Within the network of interest, every possible combination of seed-target ROIs was considered, thus leading to a 224 x 224 regression coefficients (*β*s) matrix for each participant and condition. For group-level analysis, individual *β*s were converted to *z*-scores with Fisher’s transformation. Functional contrasts, testing coupling parameters associated with one experimental condition *vs.* the other were then fed in a second level random-effect analysis. To identify conditions of interest, we applied Spatial Pairwise Clustering correction (Zalesky *et al*., 2012), as implemented in the CONN 21 toolbox. This approach is reminiscent to the cluster-based correction for multiple comparisons for fMRI activation: in this case regression coefficients are clustered together based on hierarchical cluster analysis based on both functional and spatial criteria (weighting factor 0.5). The cluster mass of connections exceeding a height threshold corresponding to *p* < 0.01 (uncorrected) was validated statistically based on the null distribution of obtained through 1000 permutations of the original dataset. In particular, clusters were identified as significant if exceeding FDR correction at p < 0.05.

**Table 1.** Stroop Task: parametric modulation of Reaction Times. The implicated regions survive FWE correction for multiple comparisons at the cluster level, with an underlying voxel-level threshold corresponding to p < 0.001. L and R refer to the left and right hemisphere, respectively. M refers to medial activations.

| | SIDE | Coordinates | | | $T_{(28)}$ | Cluster size |
| --- | --- | --- | --- | --- | --- | --- |
|  |  | x | y | z |  |  |
| Parametric modulation of Response Times |  |  |  |  |  |  |
| Middle Cingulate Gyrus | M | -10 | 20 | 30 | 7.63 | 73685*** |
| Supplementary Motor Area | M | -4 | 8 | 50 | 8.78 |  |
| Precuneus | M | -10 | -66 | 54 | 9.04 |  |
| Posterior Parietal Cortex | L | -38 | -32 | 50 | 9.55 |  |
| Posterior Parietal Cortex | R | 44 | -36 | 42 | 8.71 |  |
| Anterior Insula | L | -28 | 22 | 2 | 10.04 |  |
| Inferior Frontal Gyrus | L | -50 | 12 | 2 | 6.75 |  |
| Caudate | L | -10 | 4 | 10 | 7.66 |  |
| Thalamus | L | -10 | -18 | 8 | 8.27 |  |
| Anterior Insula | R | 30 | 26 | 0 | 8.49 |  |
| Inferior Frontal Gyrus | R | 52 | 12 | 2 | 6.07 |  |
| Caudate | R | 10 | 6 | 6 | 6.68 |  |
| Thalamus | R | 10 | -10 | 6 | 9.71 |  |
| Dorsolateral Prefrontal Cortex | R | 40 | 36 | 18 | 7.18 |  |
| Inferior Temporal Gyrus | L | -52 | -62 | -8 | 7.85 |  |
| Inferior Temporal Gyrus | R | 56 | -52 | -10 | 7.24 |  |
| Inferior Occipital Gyrus | L | -32 | -86 | 0 | 6.22 |  |
| Inferior Occipital Gyrus | R | 36 | -88 | -2 | 6.28 |  |
| Midbrain | M | 8 | -22 | -12 | 5.78 |  |
| Cerebellum | L | -24 | -70 | -48 | 7.65 |  |
| Cerebellum | R | 22 | -48 | -26 | 12.26 |  |

\*\*\* $p < 0.001$ ; corrected for multiple comparisons at the cluster level for the whole brain.

##### Multivariate Pattern Analysis

We used multivariate models predictive of subjective pain (Sharvit *et al*., 2020) and Stroop demands (Silvestrini *et al*., 2020) from neural activity, to assess whether brain-based estimates of pain unpleasantness and Stroop effect, respectively, changed as function of the manipulated conditions. The pain model is a radial-basis function kernel Support Vector Regression trained on principal components of brain activity evoked by thermal stimulations at three levels of unpleasantness from previous data. The model was subsequently validated both within the sample used for the estimation (through leave-one-out cross-validation) and on an independent cohort (see Sharvit et al., 2020). The Stroop model is a linear Support Vector Classification discriminating between Stroop Interference vs. Neutral conditions on a previous dataset (Silvestrini *et al*., 2020). Also this model was validated within the sample used for the estimation. Furthermore, we now report in Supplementary Information an independent validation on the dataset from Verstynen (2014), revealing how the original model was able to efficiently detect high Stroop demands both in terms of task preordained conditions (Interference vs. Neutral), and when modeling trials as function of participants’ Response Times though an *ad hoc* GLM where trials with longest responses in each session were specified separately from those with the most rapid reactions (through median split).

We obtained brain-based estimations of Pain Unpleasantness and Stroop demands for each participant and condition, by calculating the dot product between the model parameters and first-level brain activity. The resulting values were then analysed at the group level through the same ANOVA and t-test approaches used for the behavioural measures. Full details on how the models were developed and validated (including links to codes for their generalization to new data) are available in Supplementary Information.

## Results

### Behavioural Responses

#### Stroop performance

We confirmed that participants were both less accurate (95% *vs*. 94%, *t*_(28)_ = −2.39, *p* = 0.024, Cohen’s *d* = −0.44) and slower at responding correctly (607.98 *vs*. 628.03 ms, *t*_(28)_ = 4.47, *p* < 0.001, *d* = 0.83) during the Interference, as opposed to the Neutral condition (see Figure 2A-B). Additionally, follow up analyses revealed that task performance was negatively impacted by the onset of the thermal stimulation (see Supplementary Results for more details).

**Figure 2.**
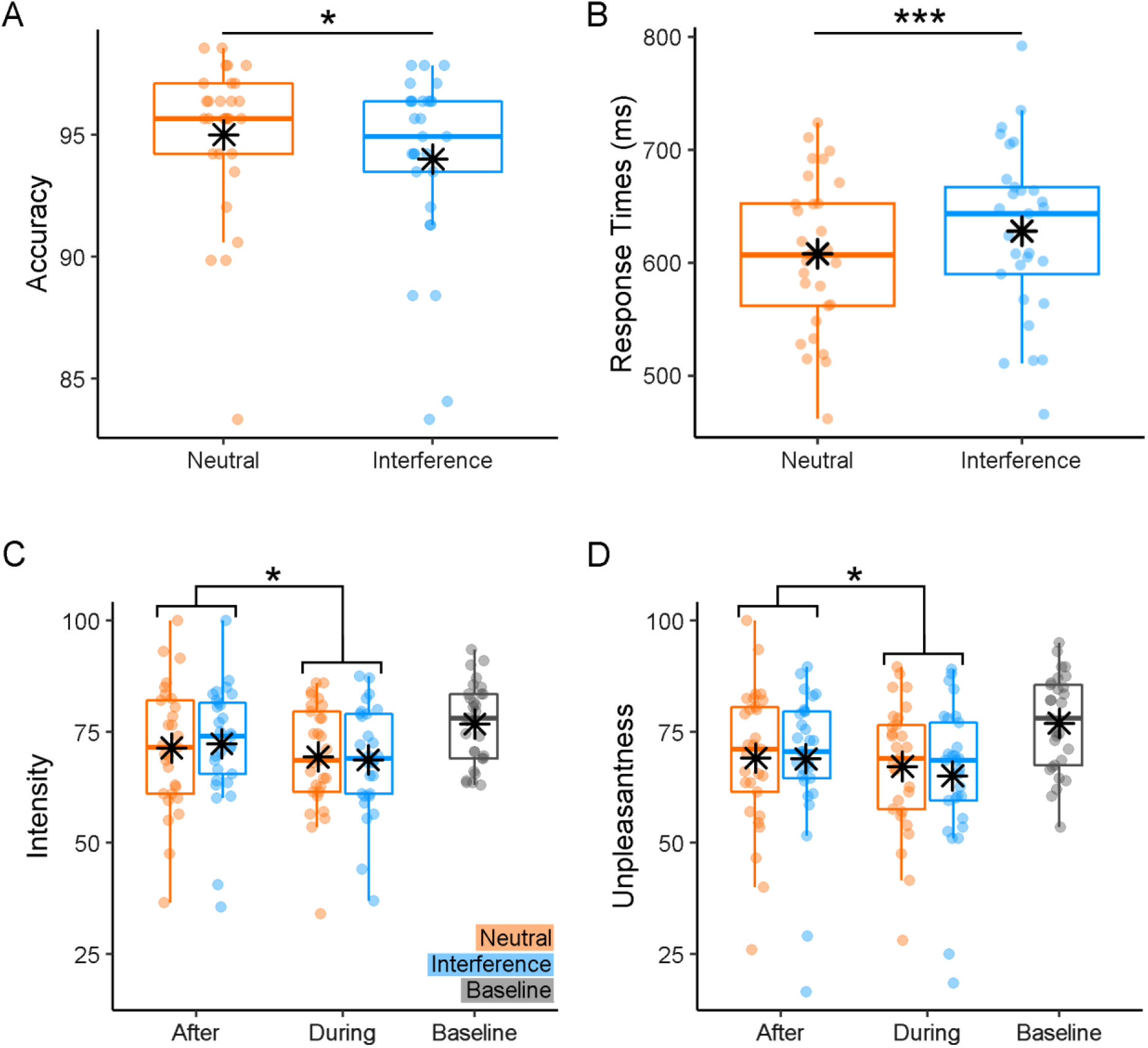
Behavioural responses. Boxplots and individual data describing (**A-B**) average Accuracy and median correct Reaction Times associated with the Stroop Task, and (**C-D**) average Pain Intensity and Unpleasantness ratings associated with the thermal stimulations. For each boxplot, the horizontal line represents the median value of the distribution, the star represents the average, the box edges refer to the inter-quartile range, and the whiskers to the data range within 1.5 of the inter-quartile range. Individual data-points are also displayed as dots. Azure boxes and dots refer to the Stroop Interference condition (and associated thermal stimulations), whereas orange boxes/dots refer to the Stroop Neutral condition, and grey boxes/dots refer to the Baseline stimulation introducing each experimental session. “***” and “*”refer to significant condition differences at *p* < 0.001 and *p* < 0.05 respectively.

#### Thermal Ratings

We analysed thermal ratings throucain effect of TIMING (Pain Intensity: *F*_(1,28)_ = 7.88, *p* = 0.014, η𝑝^2^ = 0.22; Unpleasantness: *F*_(1,28)_ = 6.25, *p* = 0.018, η𝑝^2^ = 0.19). Individuals felt painful temperatures as less intense (69.02 vs. 71.84) and unpleasant (66.17 *vs*. 69.01) *During* the task, rather *After* its completion (see Figure 2C-D). No significant modulation of TASK was found, neither as main effect nor in interaction with TIMING (*F*_(1,28)_ ≤ 1.40, *p* ≥ 0.246). Overall, our data confirm previous evidence of distraction hypoalgesia, whereby individuals consider pain less intense and unpleasant while actively engaged in a task.

### Neural Responses

#### Stroop performance

We ran a linear regression searching for regions whose activity increased monotonically with the Reaction Times. Consistently with previous studies employing Stroop (Laird *et al*., 2005; Hung *et al*., 2018), and other tasks involving cognitive control (Shackman *et al*., 2011; Hung *et al*., 2018; Kragel *et al*., 2018), we implicated a wide network involving the middle cingulate cortex, supplementary motor area, precuneus, anterior insula (AI), dorsolateral prefrontal cortex (DLPFC), posterior parietal cortex (PPC), thalamus, etc (see Table 1 and Figure 3A). To gather a more stringent functional interpretability of this network, we took a model-based approach, and analysed our data through the lens of a multivariate predictive model of Stroop high (*vs*. low) demand developed on an independent dataset (Silvestrini *et al*., 2020; see also Supplementary Information). We confirmed that this model led to a much stronger output for Stroop trials with longer Reaction Times, than those processed rapidly (*t*_(28)_ = 5.07, *p* < 0.001, *d* = 0.94; see Figure 3B). Instead, when feeding the same data to whole-brain models predictive of pain unpleasantness (Sharvit *et al*., 2020) we found no differential output between long vs. short Reaction Times in Stroop (*t*_(28)_ = 0.69, *p* = 0.497, *d* = 0.13; see Figure 3C). Overall, the activity patterns evoked by Stroop demands appear to be similar to that of previous tasks testing cognitive control via Stroop (Silvestrini *et al*., 2020), while at the same time they differ from those implicated in thermal pain (Sharvit *et al*., 2020).

**Figure 3.**
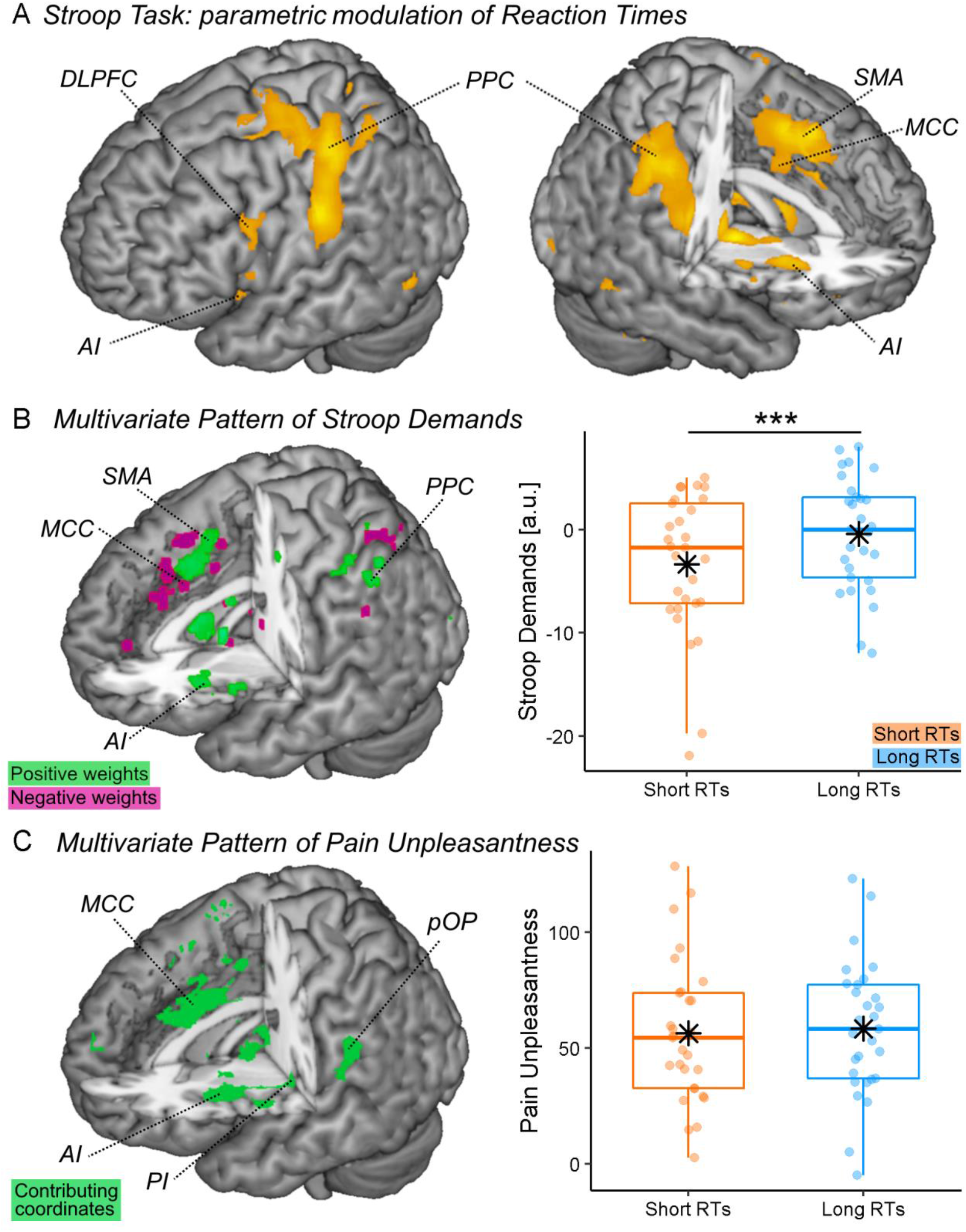
Stroop activity. (**A**) Surface rendering showing regions whose activity during the Stroop task increased linearly with the Reaction Times. To improve the readability of this specific plot, regions are displayed at voxel-level FWE correction for multiple comparisons for the whole brain, thus highlighting only brain portions showing the strongest association. (**B-C**) Predictions from whole brain models of Stroop demands (Silvestrini *et al*., 2020) and pain unpleasantness (Sharvit *et al*., 2020). Each model is described in terms of whole-brain maps with coordinates highlighted based on their relative importance. For Stroop model, coordinates describe the relative linear positive (green) or negative (violet) contribution to the prediction. For pain unpleasantness, coordinates are coded exclusively in terms of non-linear contribution (Sharvit *et al*., 2020). The output of each model is depicted separately for Stroop trials with long vs. short response times, as displayed through dedicated boxplots. Azure boxes and dots refer to trials with long response times, whereas orange boxes/dots refer to trials with short response times. Vertical values associated with Stroop Demands (in arbitrary units) reflect the degree of similarity between our data and the pattern expressed by the model (see methods). Vertical values associated with Pain unpleasantness are coded in an unpleasantness scale ranging from 0 to 100. *DLPFC*: Dorsolateral Prefrontal Cortex; *AI & PI*: Anterior & Posterior Insula; *SMA*: Supplementary Motor Area; *MCC*: Middle Cingulate Cortex; *PPC*: Posterior Parietal Cortex; *pOP*: Parietal Operculum; a.u.: arbitrary units. “***” significant condition differences at *p* < 0.001.

#### Baseline Pain

Subsequently, we inspected the neural responses associated with the delivery of painful temperatures. Figure 4 and Table 2 report brain-evoked activity associated with the baseline stimulation introducing each experimental session (relative to the non-specified parts of the model [implicit baseline]), and confirm a widespread network involving bilateral operculum, insula (in both posterior and anterior portions), PPC and right inferior frontal gyrus.

**Figure 4.**
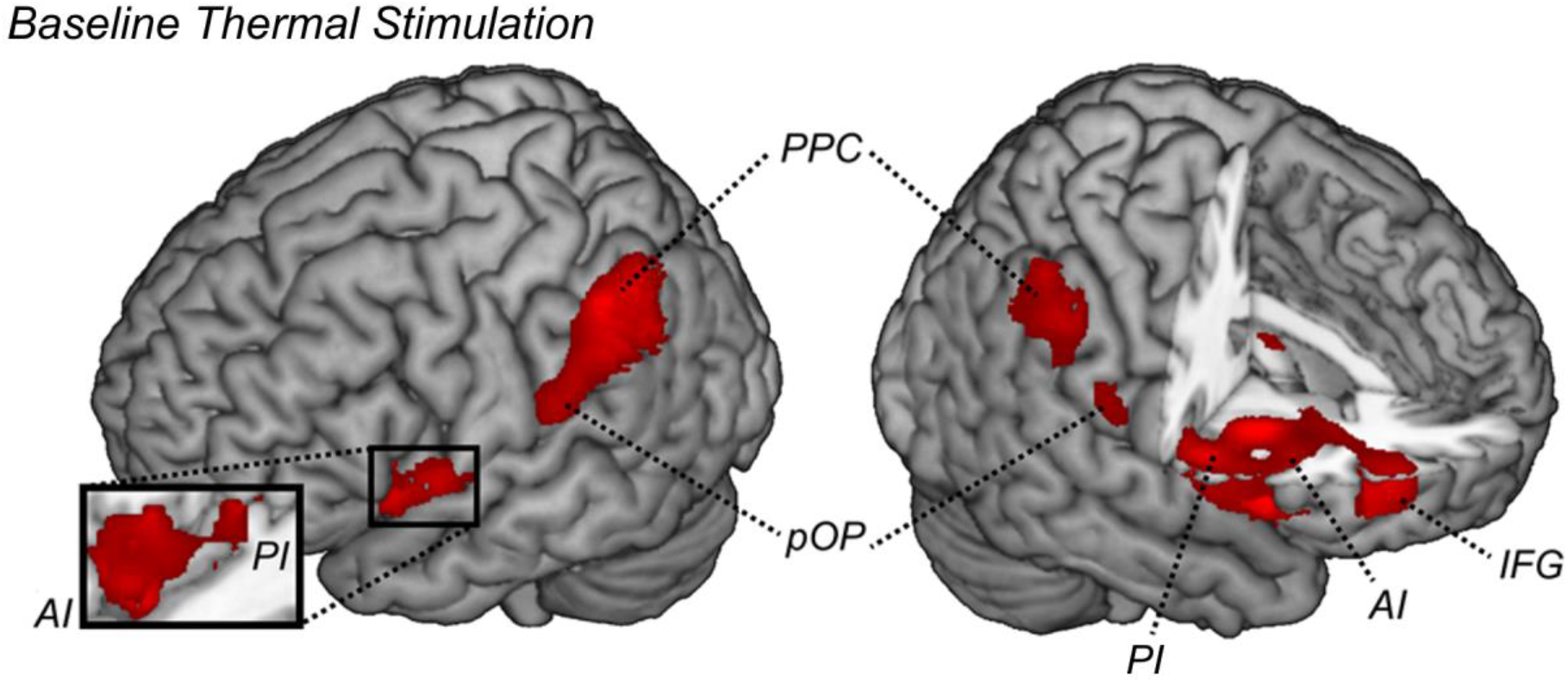
Baseline temperature. Surface rendering showing regions associated with the delivery of the baseline thermal event (when contrasted with the non-specified parts of the model [implicit baseline]). All regions displayed survive cluster-level FWE correction for multiple comparisons for the whole brain. *AI & PI*: Anterior & Posterior Insula; *pOP*: Parietal Operculum; *PPC*: Posterior Parietal Cortex; *IFG*: Inferior Frontal Gyrus.

**Table 2.**
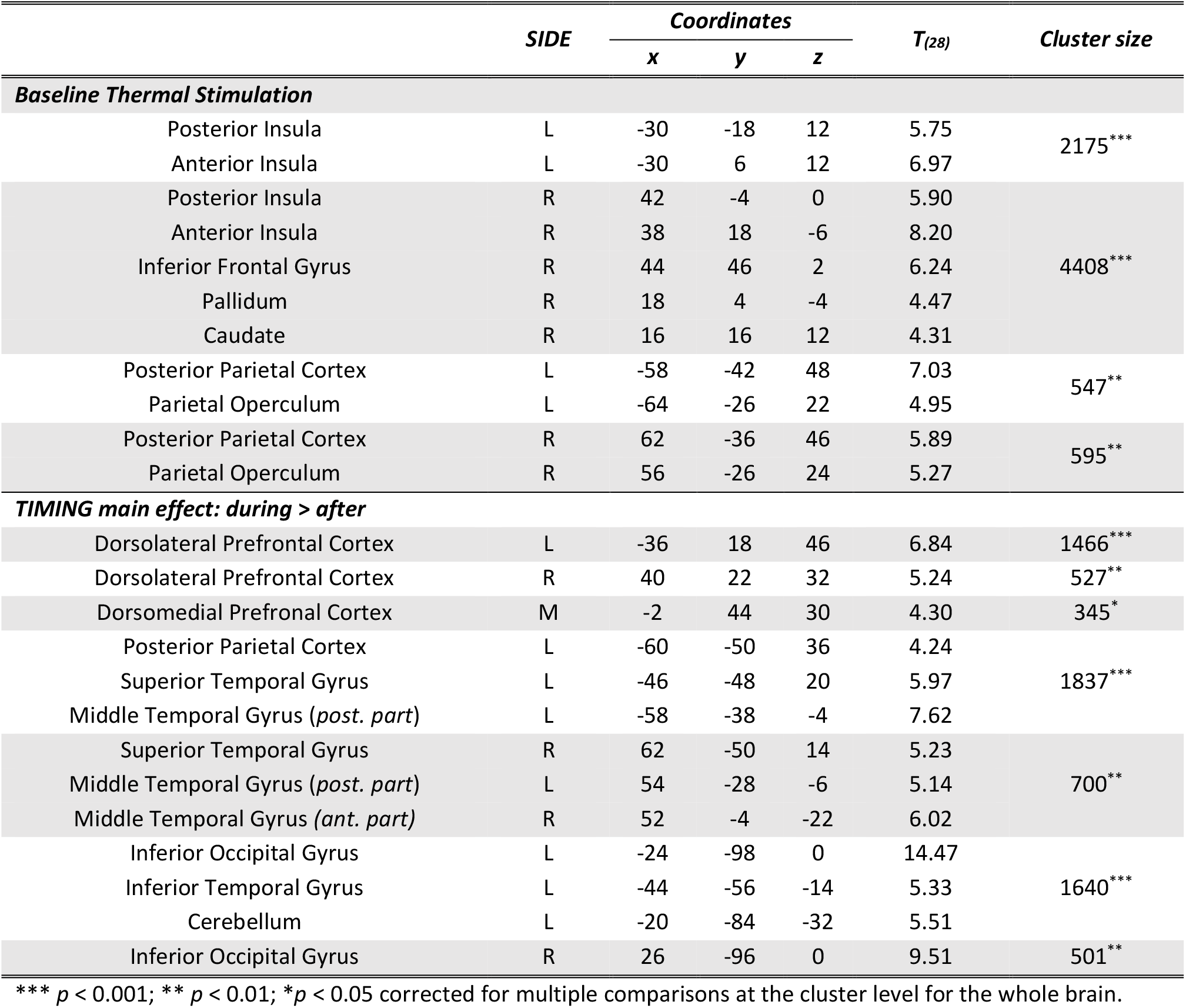
Thermal stimulations. Regions associated with the delivery of the baseline thermal event (when contrasted with the non-specified parts of the model [implicit baseline]), and with the main effect of TIMING (during > after). All regions, survive FWE correction for multiple comparisons at the cluster level, with an underlying voxel-level threshold corresponding to p < 0.001.

**Table 3.** ROI-to-ROI connectivity analysis. Change of functional connectivity associated with the main effect of TIMING. The implicated tracks survive Spatial Pairwise Clustering for multiple comparisons (Zalesky *et al*., 2012). Each cluster, is described in terms of contributing paths, t-test statistic, and global Mass index. The name of each region is drawn from the Brainnectome atlas (Fan et al., 2016, Table 1).

| Connection | $T_{(28)}$ | Mass |
| --- | --- | --- |
| *R IFJ* → *R SPL A7r* | -4.78*** | 184.31 |
| *R SPL A7c* → *R STG A38l* | -4.65*** |  |
| *R SPL A7r* → *R IFG A44d* | -4.40*** |  |
| *R SPL A7r* → *R IFJ* | -3.83*** |  |
| *R IFG A44d* → *R SPL A5l* | -3.58*** |  |
| *R SPL A5l* → *R IFG A44d* | -3.56** |  |
| *R STG A38l* → *R SPL A7c* | -3.54** |  |
| *R SPL A7c* → *R IFG A44d* | -3.49** |  |
| *R SPL A7r* → *R STG A38l* | -3.39** |  |
| *R IFG A44d* → *R SPL A7r* | -3.39** |  |
| *R SPL A7r* → *R vAI* | -3.01** |  |
| *R SPL A7c* → *R IFJ* | -2.80** |  |
| *R IFG A44d* → *R SPL A7c* | -2.80** |  |
| *R IFJ* → *R SPL A7c* | -2.80** |  |
| *L SPL A7r* → *R IFG A44d* | -5.05*** | 154.59 |
| *R IFG A44d* → *L SPL A7r* | -3.74*** |  |
| *L SPL A7r* → *R IFJ* | -3.56** |  |
| *L SPL A5l* → *R IFJ* | -3.41** |  |
| *R IFG A44d* → *L SPL A5l* | -3.35** |  |
| *R IFG A44d* → *L SPL A7ip* | -3.30** |  |
| *L SPL A5l* → *R IFG A44d* | -3.31** |  |
| *L SPL A7ip* → *R IFG A44d* | -3.27** |  |
| *L SPL A7c* → *R IFJ* | -3.21** |  |
| *L SPL A7c* → *R IFG A44d* | -3.18** |  |
| *R IFJ* → *R IFG A44d* | -3.10** |  |
| *R IFJ* → *L SPL A7c* | -2.93** |  |
| *R IFG A44d* → *L SPL A7c* | -2.27** |  |
| *R IFG A44d* → *L ACC A32p* | 4.07*** | 147.84 |
| *L ACC A32p* → *R IFG A44d* | 3.98*** |  |
| *R IFJ* → *L ACC A32sg* | 3.88*** |  |
| *L ACC A32p* → *R IFJ* | 3.86*** |  |
| *R IFG A44d* → *L ACC A32sg* | 3.45** |  |
| *R IFJ* → *L ACC A32p* | 3.43** |  |
| *R IFJ* → *R ACC A24rv* | 3.30** |  |
| *R IFJ* → *R ACC A32sg* | 3.30** |  |
| *R IFG A45c* → *L ACC A32p* | 3.26** |  |
| *L ACC A32p* → *R IFG A45c* | 3.25** |  |
| *R IFG A45c* → *L ACC A32sg* | 3.19** |  |
| *R ACC A32sg* → *R IFJ* | 2.98** |  |

#### Stroop-related Pain

##### Effects of distraction

We then tested the degree to which pain-evoked activity was influenced by the Stroop task. When analyzing the main effect of TIMING, we found stronger activity *During* (*vs. After*) the Stroop task in the bilateral DLPFC, bilateral PPC and middle temporal gyrus. Further activity was observed at the level of the dorsomedial prefrontal cortex (Figure 5A and Table 2). No region displayed stronger activity in the *After* (vs. *During*) condition. We analyzed the effect of TIMING also from a connectivity point of view. This analysis identified three clusters of connections similarly impacted by the presence of the Stroop task. In particular, the right and left PPC exhibited decreased connectivity with right DLPFC and AI (Figure 5A, green tracks). Additionally, the right DLPFC showed increased connectivity with the cingulate cortex, in both its’ dorsal (dACC) and subgenual aspects (Figure 5A, purple tracks). Please note that, although in the During condition pain stimulations and Stroop trials co-occurred, Stroop effects were independently modelled in the first-level analysis by a dedicated predictor. Hence, pain and task effects on the neural signal were not confounded in our results. Indeed, all effects presented here can be interpreted exclusively in terms of pain processing, and how they are influenced by the ongoing task.

**Figure 5.**
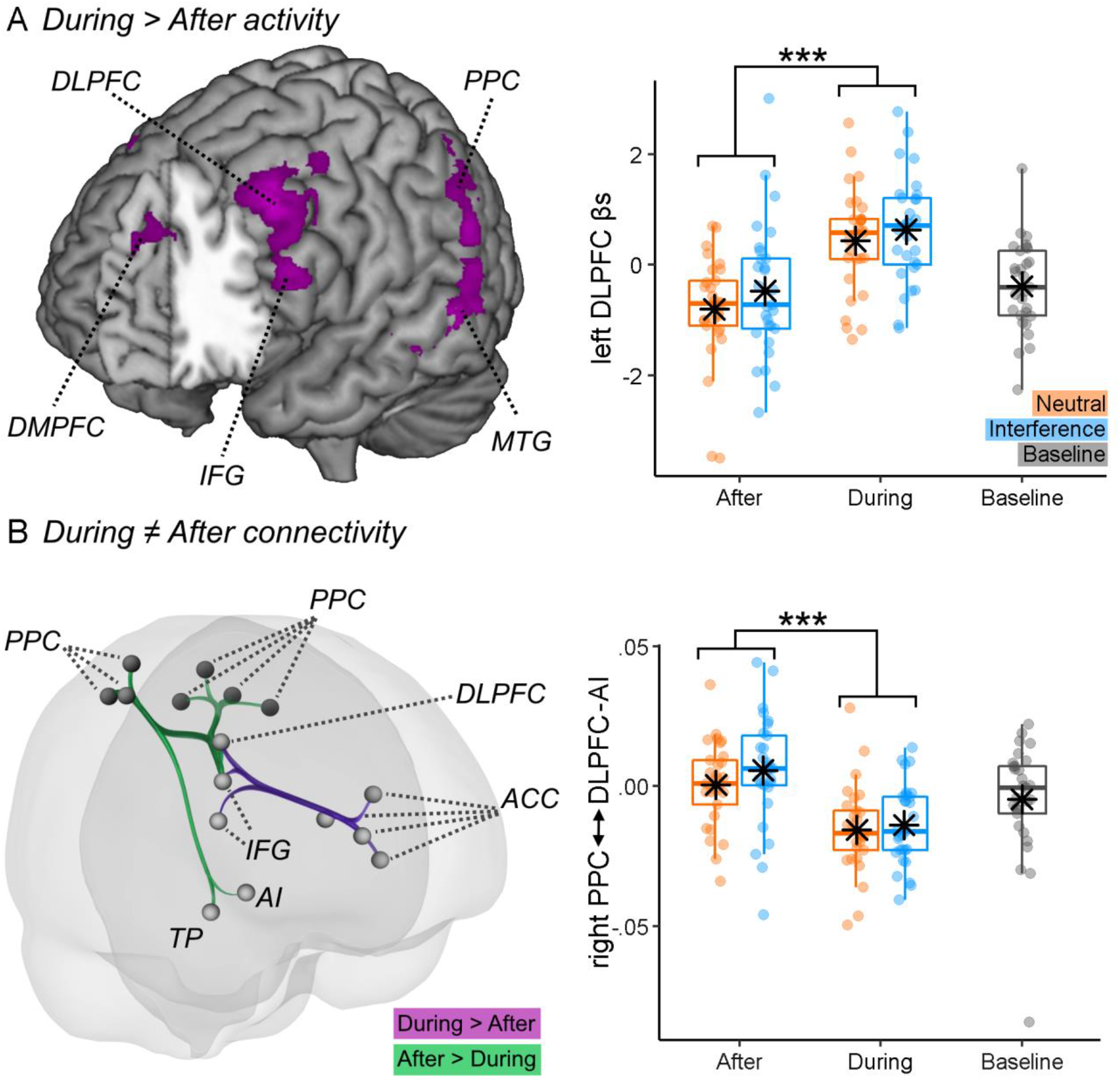
Distraction-related activity. (**A**) Surface rendering showing regions associated with the delivery of thermal stimulation *During > After* the Stroop task. All regions displayed survive FWE correction for multiple comparisons for the whole brain at *p* < 0.05. The parameter estimates from one outlined region of interest are displayed through boxplots and individual data-points. (**B**) Connectivity changes evoked by the delivery of thermal stimulations *During vs. After* the task. Purple tracks refers to increased connectivity *During > After*, whereas green tracks refer to the opposite contrast. The connectivity parameters from the track connecting right parietal cortex to lateral prefrontal cortex and anterior insula (*cluster 1*) are also displayed through boxplots and individual data-points. “***”refers to a significant main effect of TIMING at *p* < 0.001. *DLPFC & DMPFC*: Dorsolateral & Dorsomedial Prefrontal Cortex; *IFG*: Inferior Frontal Gyrus; *PPC*: Posterior Parietal Cortex; *MTG*: Middle Temporal Gyrus; *AI*: Anterior Insula; *ACC*: Anterior Cingulate Cortex; *TP*: Temporal pole.

##### Effects of Stroop demands

Subsequently, we analyzed the effect played by TASK in pain-evoked activity, both as a main effect and in interaction with TIMING. When applying correction for multiple comparisons for the whole brain no effects were observed. However, under a less conservative threshold (corresponding to *p* < 0.001 uncorrected), we found the dorsal anterior cingulate cortex (dACC) implicated in the main effect of TASK (x = 12, y = 38, z = 20, *t*_(28)_ = 4.50, 122 consecutive voxels), with stronger activity when pain was associated to the Interference *vs*. Neutral condition (Figure 6). No effect was associated with the TASK*TIMING interaction. Finally, no effects of TASK and TASK*TIMING were found at the level of connectivity.

**Figure 6.**
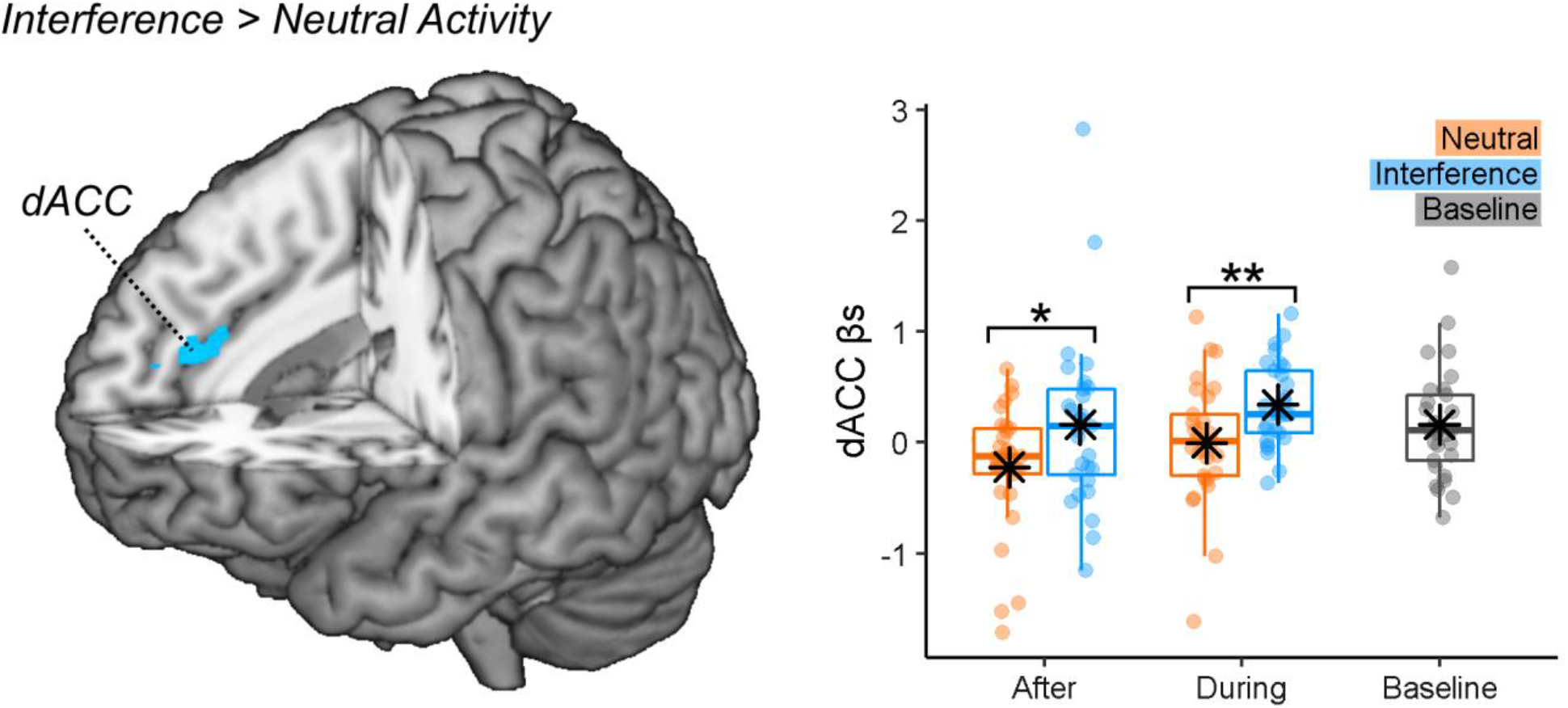
Task-demands effects on pain activity. Surface rendering showing regions associated with the delivery of thermal stimulations associated with the *Interference > Neutral* the Stroop conditions. The activations are displayed under a threshold corresponding to *p* < 0.001 (uncorrected). The parameter estimates from the outlined region are displayed through boxplots and individual data-points. “**” and “*” refer to a significant condition differences at *p* < 0.01 and *p* < 0.05 respectively. *dACC*: dorsal Anterior Cingulate Cortex.

##### Multivariate Patterns

To achieve a more reliable functional interpretation of our results, we applied to our pain-evoked responses whole-brain model predictive of pain unpleasantness (Sharvit *et al*., 2020) and Stroop demands (Silvestrini *et al*., 2020). More specifically, we analyzed the models’ output through the same ANOVA scheme used for subjective ratings. For both models we found a main effect of TIMING (Pain Unpleasantness: *F*_(1,28)_ = 8.74, *p* = 0.006, 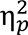 = 0.024; Stroop Demands: *F*_(1,28)_ = 5.53, *p* = 0.026, 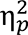 = 0.17). Figure 7 displays the models’ output across the different conditions and reveal that, consistently with what found for subjective responses, brain-estimates of Pain Unpleasantness decreased *During vs. After* the task. Instead, brain estimates of Stroop demands during pain showed an opposite trend, as they increased when participants were engaged in the task, rather than subsequent to it. No significant effects were associated with the factor TASK, neither as main effect, nor in interaction with TIMING (*F*_(1,28)_ ≤ 1.17, *p* ≥ 0.289). Finally, the effects associated with the *Pain Unpleasantness* model could be observed also when adopting a different multivariate predictive model, such as the neurological pain signature from Wager et al. (2013). Please see Supplementary Information for full details.

**Figure 7.**
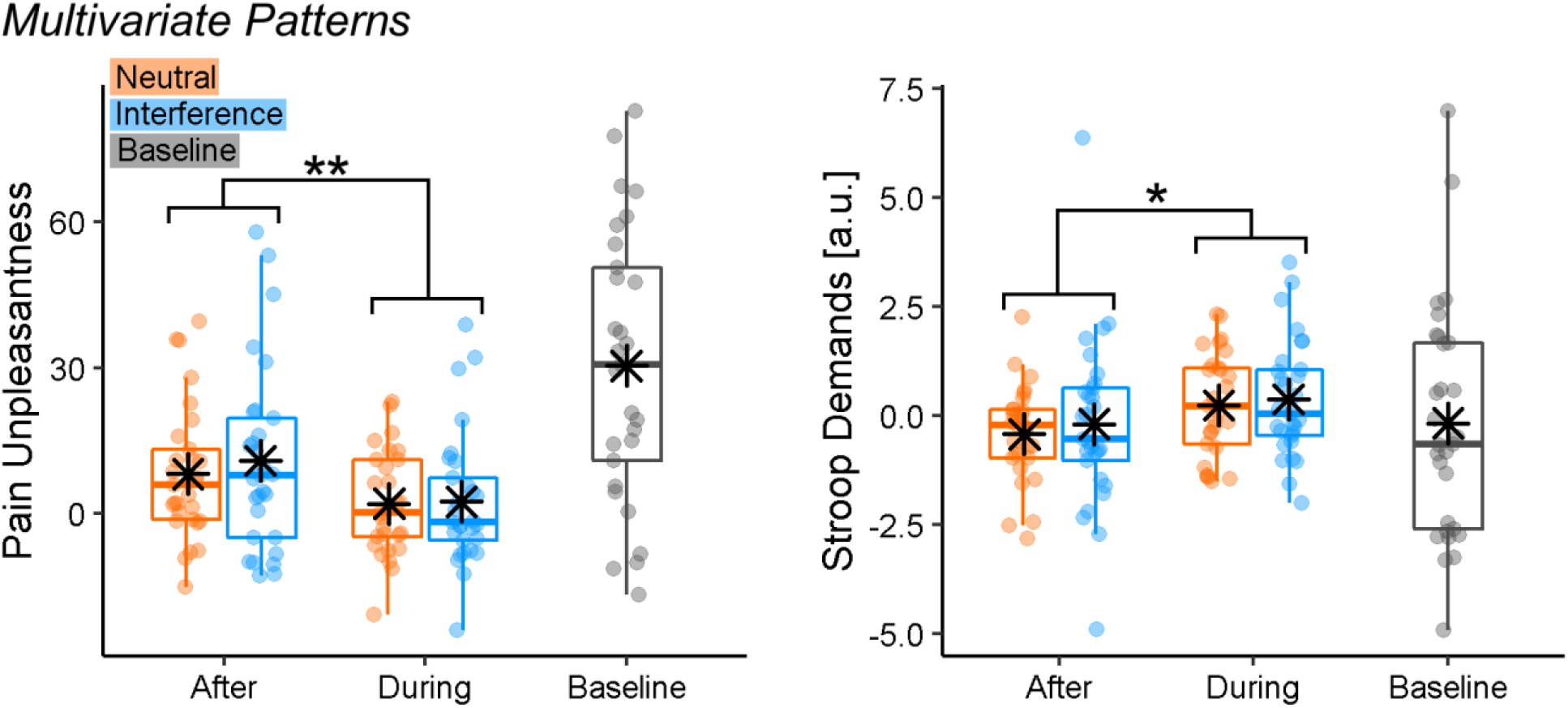
Multivariate Pattern analysis of Stroop-related pain activity. Boxplots and individual data describing the predictions from whole brain models of Stroop demands (Silvestrini *et al*., 2020) and pain unpleasantness (Sharvit *et al*., 2020), applied to the Stroop-related pain activity from our data. “**” and “*”refer to a significant main effect of TIMING from a Repeated Measures ANOVA at *p* < 0.01 and *p* < 0.05 respectively.

## Discussion

We found a dissociation in the role played by distraction and cognitive control in pain experience. Participants rated pain as less intense and unpleasant while they were engaged in an active task, consistently with previous research on distraction hypoalgesia (Petrovic *et al*., 2000; Tracey *et al*., 2002; Valet *et al*., 2004; Buhle and Wager, 2010). Such effect was associated with an increased activity at the level of prefrontal, parietal and temporal structures, but also by negative connectivity between PPC and DLPFC-AI. Finally, multivariate pattern analysis revealed that distraction altered the neural pain response, by making it less similar to that usually observed during thermal stimulations (Sharvit *et al*., 2020), and more aligned to that associated with Stroop (Silvestrini *et al*., 2020). Critically, all these effects were observed independently of cognitive control demands of the TASK employed which, instead, led to a selective increase of neural activity around dACC.

### Distraction and pain response

Our findings nicely dovetail the predictions from the neurocognitive model of attention to pain (Legrain *et al*., 2009), which speculates that PPC is associated with individual attentional set, including maintaining the focus on goal-relevant stimuli while inhibiting information from competing sources of information. Consistently, we found increased PPC activity *During* task execution (Figure 5A), and its’ negative coupling with AI (Figure 5B). This possibly reflects an inhibitory mechanism which, in turn, can explain previous evidence of decreased insular activity associated with distraction hypoalgesia (Petrovic *et al*., 2000; Valet *et al*., 2004; Seminowicz and Davis, 2007b). In this view, multiple studies showed how AI plays a key role in pain appraisal, by integrating bottom-up nociceptive signals with top-down factors related to attention and expectation (Atlas *et al*., 2010; Geuter *et al*., 2017; Sharvit *et al*., 2018). It is therefore likely that the PPC-AI inhibition observed here might nullify these top-down influences, by making one’s sensitivity more dependent on incoming ascending inputs.

In line with previous research, we also found that distraction increased activity at the level of the lateral and medial prefrontal cortex (Petrovic *et al*., 2000; Valet *et al*., 2004). However, connectivity analysis suggests that DLPFC activity is highly coupled with that of AI, but negatively associated with PPC. It is possible that, differently from PPC, DLPFC underlies compensatory processes which are engaged whenever the signal in structures like AI is high. This interpretation fits the neurocognitive model of attention to pain which speculates that DLPFC is implicated in maintaining attentional load towards the main task (Legrain *et al*., 2009), a process that could be recruited in larger extent the more competing pain signals are strong.

As for the medial portions of the prefrontal cortex, this region could play a key role triggering descending regulatory mechanisms for pain at the level of striatum, periaqueductal grey and spinal cord (Valet *et al*., 2004; Sprenger *et al*., 2015; Woo *et al*., 2015; Tinnermann *et al*., 2017). These pathways have been partly observed in previous research on distraction (Petrovic *et al*., 2000; Tracey *et al*., 2002; Valet *et al*., 2004), thus associating hypoalgesia to a well-known neurobiological model of pain regulation. In principle, the same processes could be at play also in our research, although we found no evidence that medial prefrontal cortex was interacting with the midbrain during distraction.

### Cognitive Control and pain response

When analysing the effects of TASK we found enhanced pain responses in dACC associated with an interfering Stroop (vs. neutral control), regardless of whether this condition preceded or was concurrent to the pain stimulation (Figure 6). These results fit previous findings who identify dACC as a key hub for the interplay between cognitive control and pain (Shackman *et al*., 2011; Silvestrini *et al*., 2020). Importantly, although effects of cognitive control are orthogonal to those associated with distraction, they might not be independent from them as dACC exhibits also enhanced connectivity with DLPFC-AI during task execution. Hence, pain responses in dACC are also influenced by the attentional manipulation, possibly reflecting some degree of overlap between these two facets of human executive functions.

Behaviourally we found no effect of Stroop-induced cognitive control on participants’ pain sensitivity. This might appear at odds with the literature who found significant modulations, albeit with different directions (Silvestrini and Rainville, 2013; Hoegh *et al*., 2019; Silvestrini *et al*., 2020). It is reasonable that hidden moderators/confounds might have influenced previous and current results. For instance, we recently found that cognitive control after-effects in pain are influenced by the intensity of the noxious input, with mild stimulations associated with hyperalgesia but more intense events with hypoalgesia (Riontino *et al*., 2022). Hence, it is possible that the stimulation adopted here fell within a “medium” range where its’ susceptibility to cognitive control manipulations was minimum. Future studies will need to further investigate the effect of cognitive control and pain sensitivity.

### Limitations of the study and conclusive remarks

Given the nature of the research question, our experimental design was constrained by long task sessions (Silvestrini and Rainville, 2013; Silvestrini *et al*., 2020), each associated with one thermal stimulation. This made the task-related signal susceptible to low-frequency fluctuations (which was accounted for in the analysis), whereas pain-related activity was based on limited repetitions, and with no room for a control painless stimulation. Finally, Figure 2C-D shows a strong desensitization between the first (baseline) thermal stimulation and the subsequent ones. We believe that these limitations have little impact on the main conclusions of the study, which proved to be sufficiently sensitive to capture strong thermal-related increases in the neural signal, and associated attentional effects. Furthermore, the neural modulations associated with TIME mirrored closely behavioural measures of pain and unpleasantness. As for the TASK manipulation, the lack of positive behavioural effects might reflect its poor effectiveness at influencing pain response, although the same design was effectively used before to document positive influences of Stroop on pain (Silvestrini and Rainville, 2013; Silvestrini *et al*., 2020) and vice versa (Silvestrini and Corradi-Dell’Acqua, 2022).

Finally, we acknowledge the difficulty of pulling apart entirely effects of endogenous attention and cognitive control on pain, as these two processes are highly interconnected and exert mutual influence. A more efficient way for future research could be modulating attention exogenously. However, according to the neurocognitive model of attention (Posner and Petersen, 1990), we may distinguish between executive (cognitive) control, related to conflict monitoring in tasks like the Stroop, and two other cognitive functions, namely alertness and orientation. Accordingly, one might explain distraction associated with the neutral condition in the present study as related to increased alertness independently from cognitive control. Alternatively, one might also consider that counting neutral words requires working memory, due to the maintenance of task goals’ representations, which could lead to distraction hypoalgesia (Legrain *et al*., 2009) and be associated with some degree of cognitive control. Future studies might help to further disentangle this issue.

Keeping these limitations aside, in this study we investigated the role of distraction and cognitive control in pain, with the former affecting subjective experience and triggering a widespread network involving PPC and DLPFC in interaction with AI, and the latter instead affecting specifically dACC. Overall, our study underscores for the first time the independent, and yet concurrent, role played by two different facets of human executive functions in the neural response to pain.

## Acknowledgments

NS was supported by an *Ambizione* grant awarded from the Swiss National Science Foundation (PZ00P1_142458). CCD was supported by a *Boursier* grant from the Swiss National Science Foundation (PP00P1_183715). This study was conducted at the Brain and Behavior Laboratory (BBL) at the University of Geneva and benefited from the support of the BBL technical staff.

