## Supplements for "Distraction and cognitive control independently impact parietal and prefrontal response to pain"

#### Multivariate Pattern Analysis

##### Stroop Demands

To maximize the comparability with our experiment, we choose the model from Silvestrini et al. (2020), derived from a dataset obtained when participants underwent a similar Stroop task than the one from the present study. The model was estimated and validated through the following steps.

###### *Feature Selection*

As first step, an inclusive masks was identified, including only coordinates that were involved with the term "Stroop" (either forward or reverse test) in Neurosynth toolbox. At the time when the model was estimated, this resulted in a mask comprising 1877 coordinates at a 3.5 × 3.5 × 3.5 mm resolution (Silvestrini *et al.*, 2020).

###### *Data Selection*

For modelling estimation, data from our prior work was selected (Silvestrini *et al.*, 2020), in which 24 participants underwent a similar design structure than the present study (only the "after" condition), with 2 minutes blocks involving either Stroop or control task, followed by nociceptive electrical stimulations. The modeling was based on 48 first-level parameters (24 participants * 2 conditions [Interference *vs*. Neutral]).

###### *Multivariate Modeling*

The 48 parameters associated with Stroop data were extracted from the Neurosynth Stroop mask defined before. The resulting data matrix (48 parameters × 1877 coordinates) was analysed through a linear Support Vector Machine (SVM) (Boser *et al.*, 1992) to discriminate between Stroop vs. neutral blocks based on brain activity (using a fixed regularization parameter C = 1). We employed a leave-one-subject-out cross-validation approach, to assess the proficiency of the algorithm to detect the Interference condition in an independent portion of subjects: e.g., a model trained on Stroop parameters in all-but-one subjects is used to predict pain unpleasantness in the remaining participant. The overall accuracy of the out-of-subject classification was 71%. The analysis was carried out suing the Spider toolbox for Matlab (http://people.kyb.tuebingen.mpg.de/spider). Finally, Support Vector Machine weights were used to create a whole brain map, where each coordinate within the Neurosynth Stroop mask was associated with a parameter reflecting the linear contribution to the overall prediction. This mask is currently available at the following repository (https://github.com/canlab/Neuroimaging_Pattern_Masks/tree/master/Multivariate_signature_patterns/2020_Silvestrini_Rainville_Pain_CogControl_interaction_aMCC) under the file "*stroop_pattern_wani_121416.nii*".

##### *Model Generalization*

To the best of our knowledge, no published data report evidence of generalization of the Stroop model from Silvestrini et al. (2020) on new cohorts. To account for this, we provide here validation by taking advantage of the fMRI study from Verstynen (2014), whose data is freely available under the OpenNeuro repository (Id code: *ds000164*). In this paradigm, 28 participants underwent a Stroop task similar to the one employed in both the present study and the one from Silvestrini et al. (2020). The main differences were the following. (1) Differently from our research, Verstynen (2014) implemented a colour-based Stroop by presenting single words whose meaning was inconsistent with the font colour in which they were typed. (2) The Stroop was organized in three (rather than two) conditions: in addition to *incongruent* and *neutral* events (corresponding to the two main conditions from our design), participants underwent also a *congruent* trials where task-relevant and irrelevant information led to the same answer. (3) The study was carried out in a fully event-related fashion, with the three conditions alternating randomly and intermingled by jittered inter-trial intervals (Verstynen, 2014). (4) The study was carried out in English (rather than French) and under different MRI systems and functional sequence.

For the purpose of the model validation we re-analysed the behavioural and neuroimaging data under similar routines than in the present study. Supplementary Figure 1A shows that incongruent events elicit both more errors and longer correct responses than each of the other two conditions ($|t_{(27)}| \geq 5.71$, $p < 0.001$, Cohen's $|d| \geq 1.09$). As for brain activity, we run a GLM were the three conditions were modelled separately as a delta function. Supplementary Figure 1B and Supplementary Table 1 describe the brain networks implicated in the contrast *Incongruent > Neutral*, revealing a distributed network involving the insular cortex, middle cingulate cortex, dorsolateral prefrontal cortex, supplementary motor area, posterior parietal cortex, etc. This network, highly reminiscent of the one from Silvestrini et al. (2020), was also observed for the contrast *Incongruent > Congruent* (Supplementary Table 1). Finally, we tested whether the SVM model developed by Silvestrini et al. (2020) was also sensitive to the incongruence manipulation in this study. Specifically, each of the 84 first-level images (28 subjects * 3 conditions) was resampled to the resolution of the SVM weights. Subsequently, estimates of Stroop demands was obtained as the dot-product of the resampled *βs* and the SVM weights. Hence, the higher the resulting values, the higher the adherence of the first-level images to the pattern expressed by the weight-map. The scripts for testing the proficiency of the stroop model on new data are available at the following repository https://github.com/canlab/Neuroimaging_Pattern_Masks/tree/master/Multivariate_signature_patterns/ under the file "*apply_all_signatures.m*" (option "*pattern expression*"). We found that the estimated Stroop demands was reliably stronger for the Interference condition, relative to each of the other two ($t_{(27)} \geq 4.55$, $p < 0.001$, Cohen's $d \geq 0.86$; see Supplementary Figure 1D, left subplot). Overall, we provide evidence that the model from Silvestrini et al. (2020) could generalize to a new cohort.

In the main text, we modelled Stroop trials not as function of the preordained conditions, but rather in terms of the Response Times associated with each trial. This was made necessary by long block structure of the main experiment, which would expose a condition-based analysis to low-frequency confounds, and would be vulnerable to SPM default filtering settings (see methods for more details). For consistency purposes, we exploited the dataset from Verstynen (2014) to test tested whether the model from Silvestrini et al. (2020) could generalize also Stroop data analysed in this way. Supplementary Figure 1C and Supplementary Table 1 describe the regions whose activity increases parametrically with the Response Times (regardless of the condition). The implicated network is very similar to the one observed when analysing the same data as function of manipulated incongruence (Supplementary Figure 1B), but also to the one from Figure 3A where parametrical effect of Response Times were applied to the main study. Most critically, Supplementary Figure 1D (right subplot) describe the output of the model from Silvestrini et al. (2020) in those trials with the longest responses *vs*. those with the most rapid reactions (as identified through median split, see methods section). Also in this case, the model output was significantly higher in longer response trials ($t_{(27)} = 2.71$, $p < 0.011$, Cohen's $d = 0.51$). Overall, the neural response associated with Stroop tasks led to similar effects regardless of whether they were analysed either as function of manipulated incongruence, or as function of Response Times, suggesting that both predictors underlie similar processes. In this view, Response Times represents a viable alternative for our main experiment, whose long block structure is unsuited for an analysis based on preordained experimental conditions.

#### Pain Unpleasantness

To maximize the comparability with our experiment, we choose the model from Sharvit et al. (2020) derived from datasets obtained with the same device for thermal stimulation and the same MRI scanner than the present study. The model was estimated and validated through the following steps.

##### *Feature Selection*

As first step, an inclusive masks was identified, including only coordinates that were preferentially associated with the term “pain” (association test) in the automated meta-analysis toolbox Neurosynth (Yarkoni *et al.*, 2011). At the time when the model was estimated, this resulted in a mask comprising 18759 coordinates at a 2 × 2 × 2 mm resolution (Sharvit *et al.*, 2020).

##### *Data Selection*

For modelling estimation, we selected the data from our prior work (Sharvit *et al.*, 2018), in which participants underwent three kinds of thermal stimulations held to elicit pain at low, medium and high unpleasantness, plus three kinds of olfactory stimulations of comparable unpleasantness aimed at eliciting three level of disgust (Sharvit *et al.*, 2018). Thermal events lasted 2 seconds (with 3 additional seconds for the temperature to reach plateau). In this study an expectancy manipulation was implemented, with cues sometimes predictive of a different unpleasantness (high vs. low) or modality (thermal vs. olfactory) with respect the subsequent stimulus. For the purpose of the modelling, the data from the 20 participants from Sharvit et al. (2018) was reanalyzed through an *ad hoc* first level GLM characterized by 2 functional runs for each subject, where conditions were described solely based on the bottom-up properties of the stimulations, regardless of the congruency/incongruency with the preceding cue. As our concern at the time was to compare pain and disgust neural response, 6 runs out of 40 (20 subjects * 2 runs) were excluded to prevent discrepancies in unpleasantness between thermal and olfactory events (Sharvit *et al.*, 2020). Overall pain data was based on 102 parameters (34 runs * 3 levels of pain).

##### *Data Reduction and Multivariate Modeling.*

We extracted the 102 parameters associated with pain data from the Neurosynth mask defined before. This led to a data matrix (102 parameters × 18759 coordinates) which underwent dimensionality reduction using principal component analysis (PCA), to condense the large number of coordinates in a limited number of components that retained ~ 99.9% of variance of the original dataset. This allowed us to reduce the 18759 coordinates of the pain matrix into 95 components. The scores estimated in the PCA were then modeled through Support Vector Regression (SVR) under radial basis kernel function. For validation purposes, we employed a leave-one-subject-out cross-validation, to assess the proficiency of the algorithm to predict pain unpleasantness in an independent portion of subjects. Forthermo, an additional nested cross-validation loop was included to identify at each fold the most suitable combination of hyper-parameters for the modelling ($C$, $\varepsilon$ & $\gamma$). As an overall measure of predictive proficiency, we calculated the mean squared error (MSE), reflecting the deviation between unpleasantness actually rated, and the one estimated from brain activity. The resulting MSE was considered to be significant if lower than the 5th percentile of the distribution of 1000 MSEs obtained by rerunning the same analysis procedure on permuted datasets. Overall, we found that the SVR could reliably predict pain unpleasantness in out-of-subjects data with lower error than chance (MSE = 173.97). The analysis was carried out using the LIBSVM 3.18 software (Chang and Lin, 2001).

*Model Generalization.*

We subsequently tested whether the model could predict pain unpleasantness in a different cohort. This was assessed by running the SVR analysis on the overall population, with only one cross-validation loop for hyper-parameter optimization, and then applying it to the data from Sharvit et al., (2020), where participants underwent thermal stimulation under the same hardware and timing than in Sharvit et al. (2018). The only differences with respect from the previous study were: (1) the presence of only two levels of pain unpleasantness (rather than three); (2) a different MRI functional sequence, as in Sharvit et al., (2020) we used a TR of 2.1 sec (as in the present study) whereas in Sharvit et al. (2018) we used a multiband sequence with a TR of 0.65 sec; (3) no expectancy manipulation was involved, but in part of the trials participants were exposed to a text-based storyboard prior to the thermal stimulation.

First-level images of 27 participants from Sharvit et al., (2020) were processed as follows. First, images were resliced to the same resolution of the data from Sharvit et al. (2018). Subsequently, for each subject and condition, we extracted the values from the 18759 coordinates from the same Neurosynth mask than the one used for model estimation. The 18759 $\beta$ scores were then condensed into 95 principal components, using the same PCA scores obtained during model estimation. Finally, the function *svmpredict* from LIBSVM software was used to transform the data from PCA space into kernel space, and estimate the associated unpleasantness. We found that the estimated pain unpleasantness was reliably stronger for high *vs*. low stimulations, regardless of whether participants were exposed to a text-based storyboard or not ($t_{(26)} \geq 4.69$, $p \leq 0.001$; see also Figure 5 from Sharvit et al., 2020, for graphical representation). The pain model, and the scripts for testing its proficiency on new data are available under the Open Science Framework at https://osf.io/jkrvp/.

Importantly, the model from Sharvit et al. (2020) mapped unpleasantness in a scale ranging from 0 (neutral) to 50 (extremely unpleasant). However, in the present study, unpleasantness was measured on a 0-100 scale. For this reason, when applying the model to our current data, we multiplied the output by two, to fit the 0-100 scale.

##### Applying brain signatures to our data.

We applied two above-described model to the neural responses from our study. The main text describes modulations at the level of Stroop-effects on pain activity, whereas here we describe follow-up results associated with the first-level parameter images from the baseline temperature. In particular, the output of the Pain Unpleasantness model correlated significantly with the stimulation temperature (Pearson's $r$ = 0.46, $p$ = 0.013; Supplementary Figure 2, left plot). Please note that the temperature was optimized at the individual level to account for individual differences in pain sensitivity, and that these differences in nociceptive intensity were reliably captured by this pain model. Interestingly, when analysing the same data under the lens of the Stroop demands model, a similar positive relationship between predicted outcome and stimulation temperature was observed ($r$ = 0.44, $p$ = 0.017; Supplementary Figure 2, left plot). Hence, although pain and cognitive control activity patterns dissociate at the neural level (Kragel *et al.*, 2018; Silvestrini *et al.*, 2020), it is possible to observe the occurrence of both in the neural response evoked by painful stimulations (Silvestrini *et al.*, 2020).

We also wished to insure that our main results associated with the Pain Unpleasantness model, were not idiosyncratic to the algorithm adopted. As such we repeated the analyses (including the ones from the main text) by using instead the seminal Neurological Pain Signature from Wager et al. (2013). This is a linear model of pain intensity developed through a PCA-LASSO algorithm and validated across multiple datasets, and across different kinds of nociceptive stimulations and acquisition parameters (Wager *et al.*, 2013; Krishnan *et al.*, 2016). The model is structured as a whole brain map, where each coordinate was associated with a parameter reflecting the linear contribution to the overall prediction. We applied this model to the pain-evoked activity in our dataset by estimating the dot-product of the *βs* images and the model parameters, using the same routines described above for the Stroop model. Results are displayed in Supplementary Figure 3, and lead to a very similar result pattern to the one described in the main text, with very similar effect sizes.

In particular, the output of the Neurological Pain Signature was modulated by the factor TIMING, revealing lower pain estimates when the thermal stimulation was delivered during (vs. after) the task ($F_{(1,28)}$ = 4.87, $p$ = 0.036, $\eta_p^2$ = 0.15; Supplementary Figure 3A). Furthermore, neither the main effect of TASK nor the TASK*TIMING interaction were found to ($F_{(1,28)} \leq 0.37$, $p \geq 0.135$). Furthermore, the output of the baseline stimulation was positively correlated with the temperature used (Pearson's $r$ = 0.38, $p$ = 0.043; Supplementary Figure 3B). Overall, this follow-up analysis shows that our main results were not idiosyncratic to the model adopted, but could be observed by using also other pain signatures.

#### Impact of Pain on Performance

Our design was not optimized for specifically analysing the impact of pain on performance because the painful stimulation only occurred during 4 trials of the task. This made it hard and probably not appropriate to detect any effect. However, for full disclosure, we provide this analysis in this supplemental material as suggested in the comments of one of the external reviewers.

In particular, for both Accuracy and correct Response Times, we run a repeated measures ANOVA focused exclusively on the portion of data associated with the *During* condition. In each of this model we tested the effect of Stroop (Interference vs. Neutral) and that of Pain (before *vs*. after the onset of the thermal stimulation). To compare Pain effect in a balanced setting, we considered the 4 trials following the onset of the thermal stimulation, and those 4 immediately preceding it. The analysis of the median Response Times, provided an effect of Pain ($F_{(1,28)}$ = 6.07, $p$ = 0.020, $\eta_p^2$ = 0.18) revealing higher Response times following the onset of pain relative to before (638.21 *vs*. 619.22 ms). No other effect was found to be significant ($F_{(1,28)} \leq 1.49$, $p$ = 0.233). Instead, the analysis of average Accuracy led to no effect whatsoever ($F_{(1,28)} \leq 2.38$, $p \leq 0.134$).

Overall, this analysis provides evidence that the onset of pain interfered on participants' performance on the Stroop (at least when Response Times are concerned). Importantly, this analysis was obtained comparing the last 8 trials of each block (4 before *vs*. 4 after pain onset), thus making unlikely any confound related on overall fatigue (which would be present when comparing the beginning vs. the end of each block). However, we should remind that the limited number of trials suggest caution in the interpretation of these results.

#### Supplementary Figures

##### Supplementary Figure 1

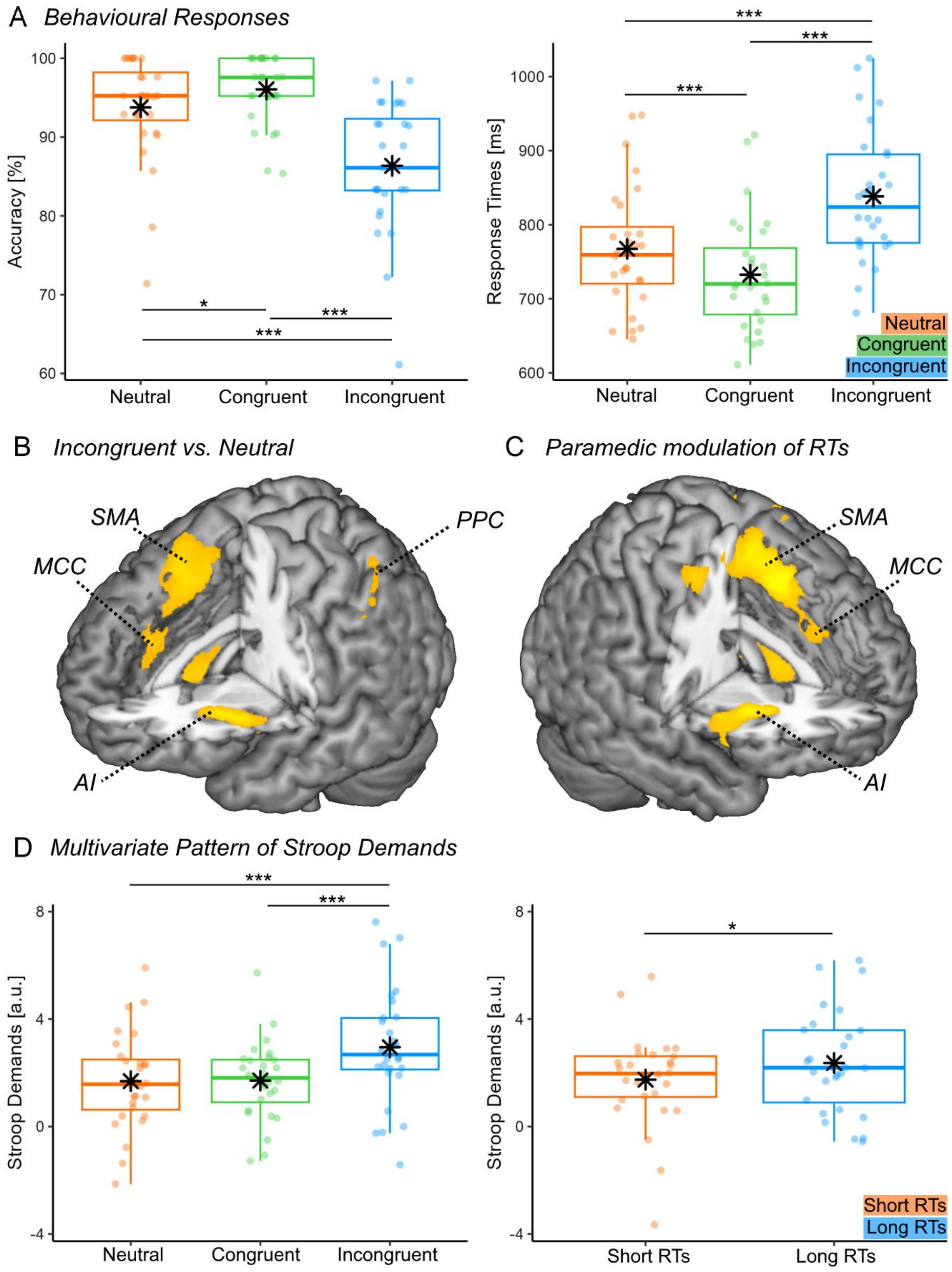


*Re-analysis of the data from Verstynen (2014). (**A**) Behavioural responses. Boxplots and individual data describing average Accuracy and correct Reaction Times associated with the Stroop Task. For each boxplot, the horizontal line represents the median value of the*

*distribution, the star represents the average, the box edges refer to the inter-quartile range, and the whiskers to the data range within 1.5 of the inter-quartile range. Individual data-points are also displayed as dots. Azure boxes and dots refer to the Stroop incongruent condition, green boxes/dots refer to the congruent condition, whereas orange boxes/dots refer to the Neutral condition. "***" and "*"refer to significant condition differences assessed through paired-sample t-test at $p < 0.001$ and $p < 0.05$ respectively. (**B-C**) Surface rendering showing regions whose activity during the Stroop task was stronger in incongruent vs. neutral events (B), or increased linearly with the Reaction Times regardless of the condition (C). All regions displayed survive cluster-level FWE correction for multiple comparisons for the whole brain. AI Anterior Insula; SMA: Supplementary Motor Area; MCC: Middle Cingulate Cortex; PPC: Posterior Parietal Cortex; pOP: Parietal Operculum. (**D**) Multivariate Pattern analysis. Boxplots and individual data describing the output from whole brain models of Stroop demands (Silvestrini et al., 2020) applied to the Stroop-related pain activity from our data. The left subplot describes the model output obtained when analysing the data in terms preordained conditions, under the same color-coding adopted for the display of behavioural results. The right subplots describes the model output obtained when analysing the data in terms of Response Times. In the right subplot only azure boxes/dots refer to trials associated with the longest Response Times, whereas orange boxes/dots refer to trials associated with the shortest Response Times.*

**Supplementary Figure 2**

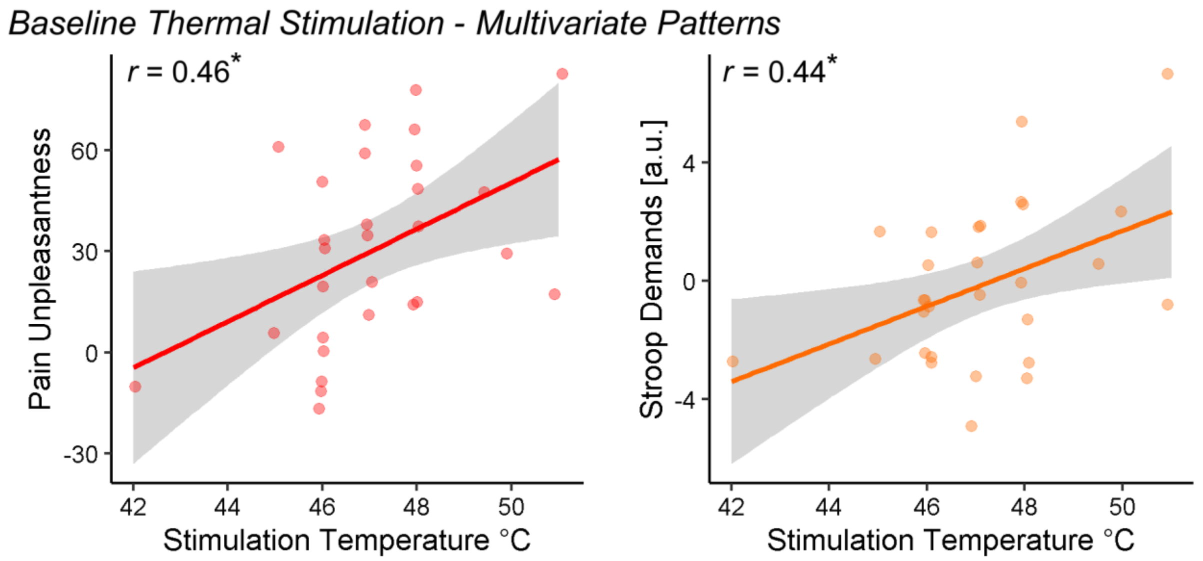


*Linear regressions (with confidence interval areas) and individual data describing the predictions from whole brain models of Stroop demands (Silvestrini et al., 2020) and pain unpleasantness (Sharvit et al., 2020) plotted against the stimulation temperature. "*" refers to significant Pearson's r coefficient at $p < 0.05$.*

**Supplementary Figure 3**

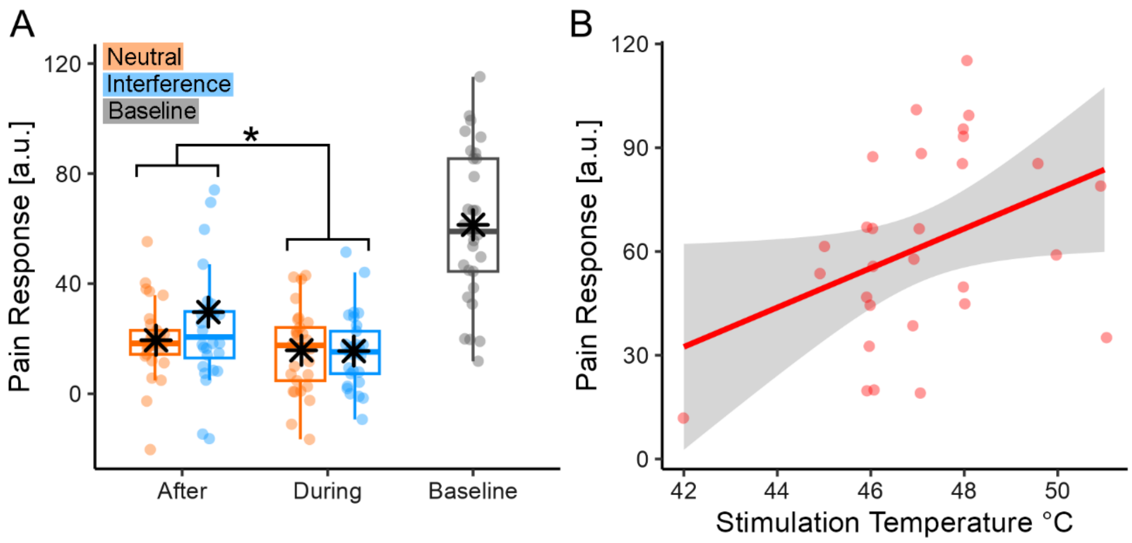


***(A)*** *Multivariate Pattern analysis of Stroop-related pain activity. Boxplots and individual data describing the predictions from Neurological Pain Signature (Wager et al., 2013), applied to the Stroop-related pain activity from our data. "*"refer to a significant main effect of TIMING from a Repeated Measures ANOVA at p < 0.01 and p < 0.05 respectively.* ***(B)*** *Baseline Temperature: linear regressions (with confidence interval areas) and individual data describing the predictions from the Neurological Pain Signature plotted against the stimulation temperature.*

### Supplementary Tables

#### Supplementary Table 1

Re-analysis of the data from Verstynen (2014). Regions implicated in the contrast Incongruent > Neutral, Incongruent > Congruent and in the contrast testing the parametric modulation of Response Times. The implicated regions survive FWE correction for multiple comparisons at the cluster level, with an underlying voxel-level threshold corresponding to *$p < 0.001$*. L and R refer to the left and right hemisphere, respectively. M refers to medial activations.

| | *SIDE* | *Coordinates* | | | $T_{(27)}$ | *Cluster size* |
|---|---|---|---|---|---|---|
| | | *x* | *y* | *z* | | |
| ***Incongruent > Neutral*** | | | | | | |
| Anterior Insula | R | 32 | 22 | 6 | 7.63 | $2278^{***}$ |
| Dorsolateral Prefrontal Cortex | R | 46 | 18 | 28 | 5.25 | |
| Anterior Insula | L | -30 | 22 | -10 | 7.58 | $2782^{***}$ |
| Dorsolateral Prefrontal Cortex | L | -34 | 0 | 50 | 6.46 | |
| Posterior Parietal Cortex | R | 38 | -44 | 36 | 4.72 | $262^{*}$ |
| Posterior Parietal Cortex | L | -30 | -52 | 44 | 5.30 | $952^{***}$ |
| Middle Cingulate Cortex | M | 6 | 32 | 26 | 4.56 | $1426^{***}$ |
| Supplementary Motor Area | M | -6 | 16 | 50 | 6.95 | |
| Thalamus | R | 10 | -6 | -8 | 5.41 | $638^{**}$ |
| Caudate | R | 12 | 14 | 0 | 5.18 | |
| Precuneus | M | -4 | -66 | 44 | 4.35 | $247^{*}$ |
| ***Incongruent > Congruent*** | | | | | | |
| Anterior Insula | R | 32 | 24 | 40 | 8.29 | $1457^{***}$ |
| Dorsolateral Prefrontal Cortex | R | 44 | 20 | 26 | 5.74 | $567^{**}$ |
| Anterior Insula | L | -30 | 22 | -10 | 5.77 | $473^{**}$ |
| Dorsolateral Prefrontal Cortex | L | -48 | 14 | 24 | 6.82 | $1238^{***}$ |
| Posterior Parietal Cortex | R | 48 | -38 | 44 | 4.01 | $249^{*}$ |
| Superior Temporal Gyrus | R | 64 | -44 | 18 | 4.97 | |
| Posterior Parietal Cortex | L | -28 | -58 | 44 | 4.72 | $859^{***}$ |
| Middle Cingulate Cortex | M | 10 | 28 | 36 | 5.23 | $999^{***}$ |
| Supplementary Motor Area | M | -8 | 8 | 58 | 4.93 | |
| Thalamus | R | 10 | -6 | -2 | 7.39 | $899^{***}$ |
| Caudate | R | 12 | 12 | -2 | 4.56 | |
| Hippocampus | R | 24 | -30 | -6 | 3.89 | |
| Brain Stem | M | 2 | -22 | -24 | 3.72 | |
| Calcarine Cortex | R | 14 | -68 | 12 | 6.24 | $404^{**}$ |
| Calcarine Cortex | L | -10 | -74 | 6 | 5.17 | $318^{*}$ |
| ***Parametrical modulation of Response Times*** | | | | | | |
| Anterior Insula | R | 32 | 18 | 2 | 7.22 | $1362^{***}$ |
| Anterior Insula | L | -28 | 26 | -2 | 5.27 | $9759^{***}$ |
| Dorsolateral Prefrontal Cortex | L | -42 | 20 | 24 | 5.33 | |
| Precentral Gyrus | L | -26 | 0 | 56 | 7.96 | |
| Posterior Parietal Cortex | L | -34 | -40 | 42 | 5.69 | |
| Supplementary Motor Area | M | -4 | 14 | 50 | 7.42 | |
| Middle Cingulate Cortex | M | 10 | 30 | 24 | 5.60 | |

| | | | | | | |
|---|---|---|---|---|---|---|
| Thalamus | L | -10 | -18 | 12 | 5.84 | 596** |
| Caudate | L | -12 | 8 | 4 | 4.96 | |

*** $p < 0.001$; ** $p < 0.01$; *$p < 0.05$ corrected for multiple comparisons at the cluster level for the whole brain.
